## Supplement_Figure_S1-S9_Table_S1-S2 for "The Kelch13 compartment is a hub of highly divergent vesicle trafficking proteins in malaria parasites"

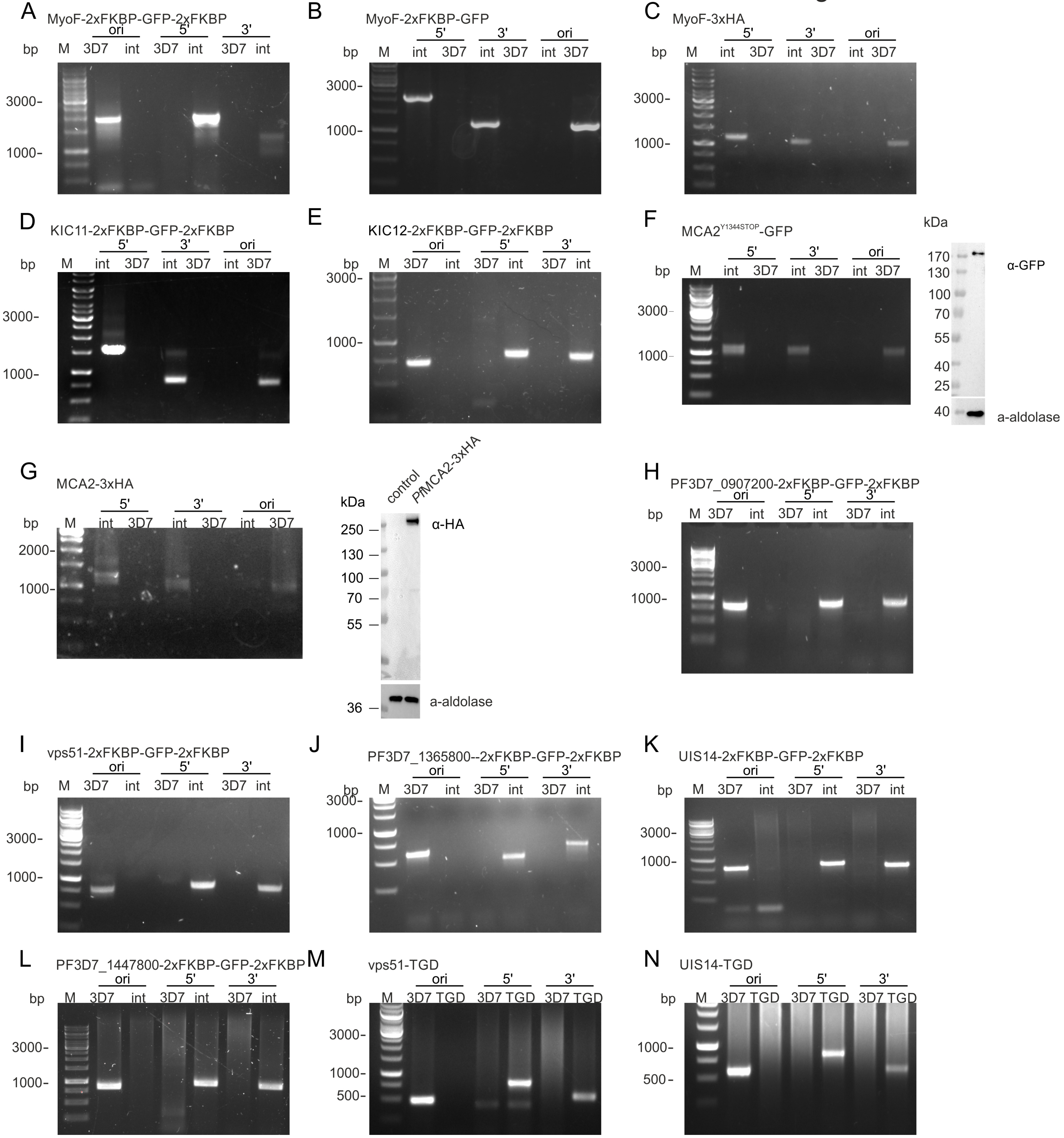
A
MyoF-2xFKBP-GFP-2xFKBP
ori
5'
3'
bp
M
3D7
int
3000
1000
B
MyoF-2xFKBP-GFP
C
MyoF-3xHA
D
KIC11-2xFKBP-GFP-2xFKBP
E
KIC12-2xFKBP-GFP-2xFKBP
F
MCA2Y1344STOP-GFP
kDa
170
130
100
70
55
40
25
α-GFP
a-aldolase
G
MCA2-3xHA
2000
control
PfMCA2-3xHA
250
36
α-HA
H
PF3D7_0907200-2xFKBP-GFP-2xFKBP
I
vps51-2xFKBP-GFP-2xFKBP
J
PF3D7_1365800--2xFKBP-GFP-2xFKBP
K
UIS14-2xFKBP-GFP-2xFKBP
L
PF3D7_1447800-2xFKBP-GFP-2xFKBP
M
vps51-TGD
TGD
500
N
UIS14-TGD

### Figure S2

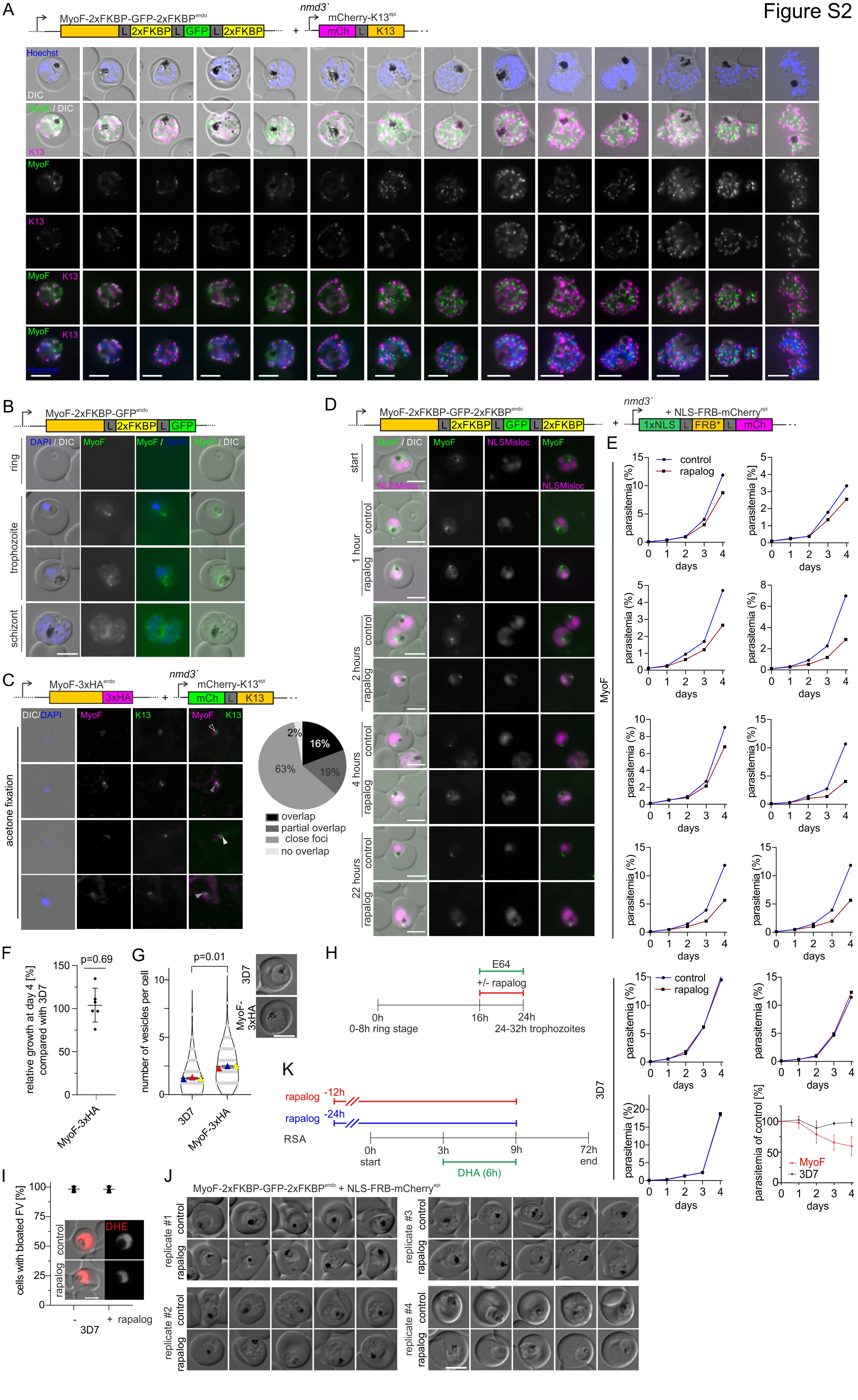
A
MyoF-2xFKBP-GFP-2xFKBP endo
L 2xFKBP L GFP L 2xFKBP
+
nmd3`
mCherry-K13 epi
mCh L K13
Hoechst
DIC
MyoF / DIC
K13
MyoF
K13
MyoF K13
MyoF K13 Hoechst
B
MyoF-2xFKBP-GFP endo
L 2xFKBP L GFP
DAPI / DIC
MyoF
MyoF / DAPI
MyoF / DIC
ring
trophozoite
schizont
C
MyoF-3xHA endo
3xHA
+
nmd3`
mCherry-K13 epi
mCh L K13
DIC/DAPI
MyoF
K13
MyoF K13
acetone fixation
2%
16%
63%
19%
overlap
partial overlap
close foci
no overlap
D
MyoF-2xFKBP-GFP-2xFKBP endo
L 2xFKBP L GFP L 2xFKBP
+
nmd3`
+ NLS-FRB-mCherry epi
1xNLS L FRB* L mCh
MyoF / DIC
MyoF
NLSMisloc
MyoF
NLSMisloc
NLSMisloc
start
1 hour
2 hours
4 hours
22 hours
control
rapalog
E
control
rapalog
parasitemia (%)
parasitemia [%]
days
MyoF
3D7
parasitemia of control [%]
MyoF
3D7
F
p=0.69
relative growth at day 4 [%] compared with 3D7
MyoF-3xHA
G
p=0.01
number of vesicles per cell
3D7
MyoF-3xHA
3D7
MyoF-3xHA
H
E64
+/- rapalog
0h
0-8h ring stage
16h
24h
24-32h trophozoites
K
rapalog -12h
rapalog -24h
RSA
0h start
3h
9h
72h end
DHA (6h)
I
cells with bloated FV [%]
control
rapalog
DHE
-
+ rapalog
3D7
J
MyoF-2xFKBP-GFP-2xFKBP endo + NLS-FRB-mCherry epi
replicate #1
replicate #2
replicate #3
replicate #4
control
rapalog

Figure S3

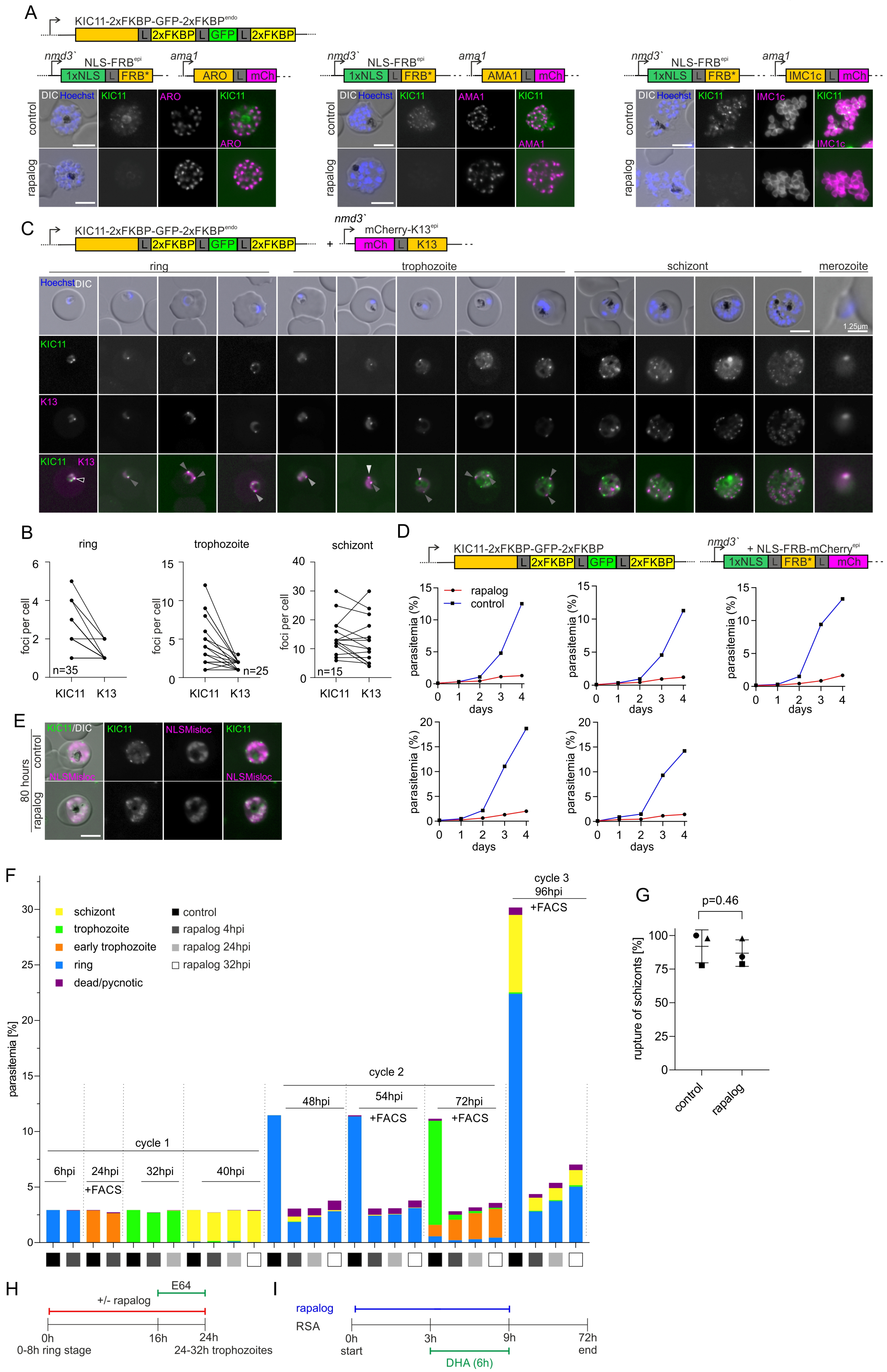

A
KIC11-2xFKBP-GFP-2xFKBP
nmd3`
NLS-FRB
ama1
1xNLS
FRB*
ARO
AMA1
IMC1c
mCh
DIC Hoechst
KIC11
control
rapalog
C
mCherry-K13
K13
ring
trophozoite
schizont
merozoite
1.25µm
B
foci per cell
n=35
n=25
n=15
D
rapalog
control
parasitemia (%)
days
E
80 hours
NLSMisloc
KIC11/DIC
F
schizont
trophozoite
early trophozoite
ring
dead/pycnotic
rapalog 4hpi
rapalog 24hpi
rapalog 32hpi
parasitemia [%]
cycle 1
cycle 2
cycle 3
6hpi
24hpi
32hpi
40hpi
48hpi
54hpi
72hpi
96hpi
+FACS
G
p=0.46
rupture of schizonts [%]
H
+/- rapalog
E64
0h
16h
24h
0-8h ring stage
24-32h trophozoites
I
RSA
3h
9h
72h
start
end
DHA (6h)

Figure S4

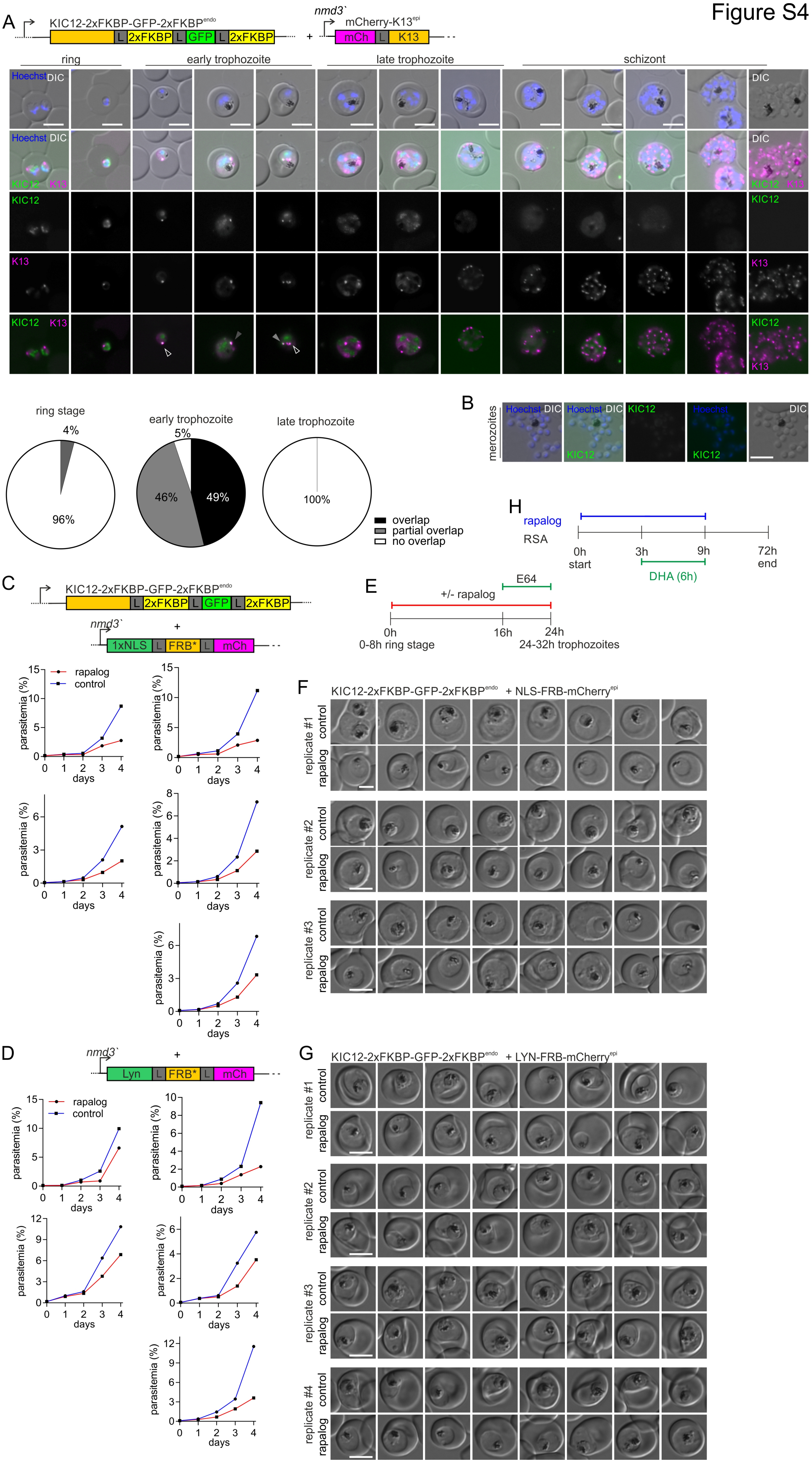

A
KIC12-2xFKBP-GFP-2xFKBP^endo
L 2xFKBP L GFP L 2xFKBP
+
nmd3`
mCherry-K13^epi
mCh L K13
ring
early trophozoite
late trophozoite
schizont
Hoechst DIC
KIC12 K13
KIC12
K13
KIC12 K13
ring stage
4%
96%
early trophozoite
5%
46%
49%
late trophozoite
100%
overlap
partial overlap
no overlap
B
merozoites
Hoechst DIC
KIC12
DIC
C
KIC12-2xFKBP-GFP-2xFKBP^endo
nmd3`
1xNLS L FRB* L mCh
rapalog
control
parasitemia (%)
days
D
Lyn L FRB* L mCh
E
+/- rapalog
E64
0h
16h
24h
0-8h ring stage
24-32h trophozoites
F
KIC12-2xFKBP-GFP-2xFKBP^endo + NLS-FRB-mCherry^epi
replicate #1
replicate #2
replicate #3
G
KIC12-2xFKBP-GFP-2xFKBP^endo + LYN-FRB-mCherry^epi
replicate #4
H
rapalog
RSA
0h
start
3h
9h
72h
end
DHA (6h)

Figure S5

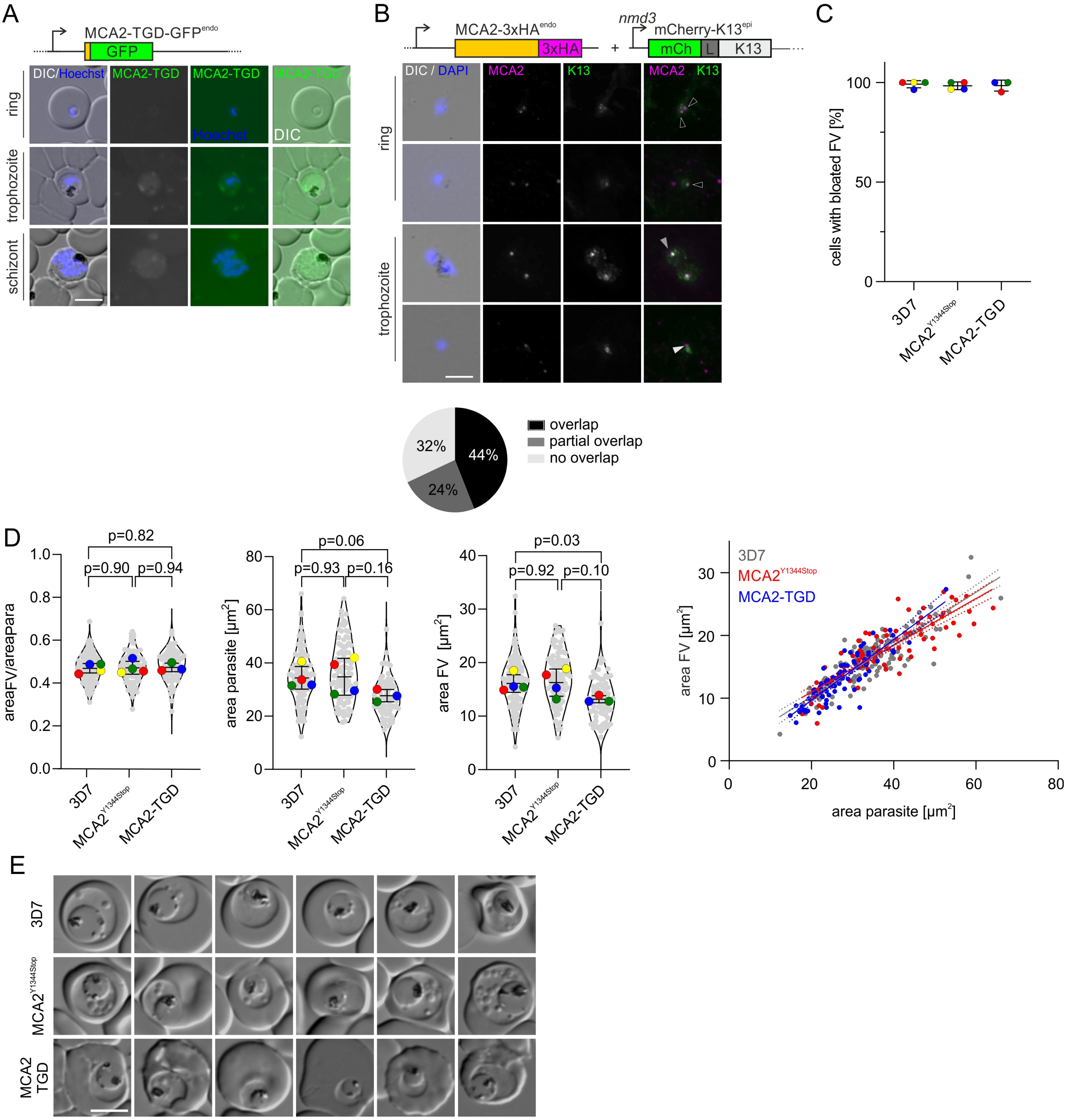

A
MCA2-TGD-GFPendo
GFP
DIC/Hoechst
MCA2-TGD
MCA2-TGD
MCA2-TGD
Hoechst
DIC
ring
trophozoite
schizont
B
MCA2-3xHAendo
3xHA
+
nmd3
mCherry-K13epi
mCh
L
K13
DIC / DAPI
MCA2
K13
MCA2 K13
ring
trophozoite
32%
44%
24%
overlap
partial overlap
no overlap
C
cells with bloated FV [%]
100
50
0
3D7
MCA2Y1344Stop
MCA2-TGD
D
p=0.82
p=0.90
p=0.94
areaFV/areapara
1.0
0.8
0.6
0.4
0.2
0.0
p=0.06
p=0.93
p=0.16
area parasite [µm2]
80
60
40
20
0
p=0.03
p=0.92
p=0.10
area FV [µm2]
40
30
20
10
0
3D7
MCA2Y1344Stop
MCA2-TGD
area FV [µm2]
30
20
10
0
0
20
40
60
80
area parasite [µm2]
E
3D7
MCA2Y1344Stop
MCA2 TGD

### Figure S6

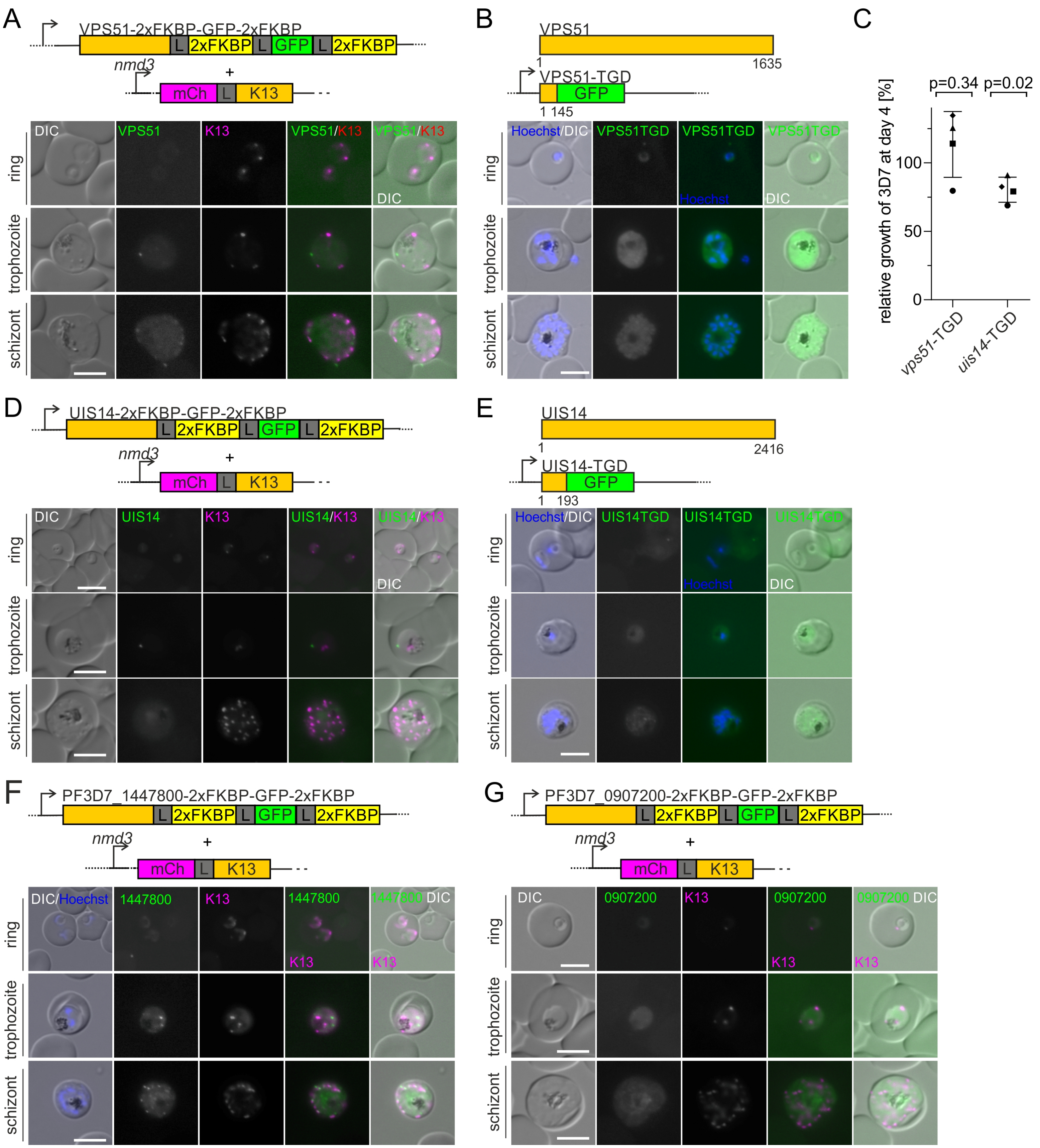
A
VPS51-2xFKBP-GFP-2xFKBP
L 2xFKBP L GFP L 2xFKBP
nmd3
+
mCh L K13
DIC
VPS51
K13
VPS51/K13
VPS51/K13 DIC
ring
trophozoite
schizont
B
VPS51
1
1635
VPS51-TGD
GFP
1 145
Hoechst/DIC
VPS51TGD
VPS51TGD Hoechst
VPS51TGD DIC
C
p=0.34
p=0.02
relative growth of 3D7 at day 4 [%]
0
50
100
vps51-TGD
uis14-TGD
D
UIS14-2xFKBP-GFP-2xFKBP
UIS14
UIS14/K13
UIS14/K13 DIC
E
UIS14
1
2416
UIS14-TGD
1 193
UIS14TGD
UIS14TGD Hoechst
UIS14TGD DIC
F
PF3D7_1447800-2xFKBP-GFP-2xFKBP
DIC/Hoechst
1447800
1447800 K13
1447800 DIC K13
G
PF3D7_0907200-2xFKBP-GFP-2xFKBP
0907200
0907200 K13
0907200 DIC K13

### Figure S7

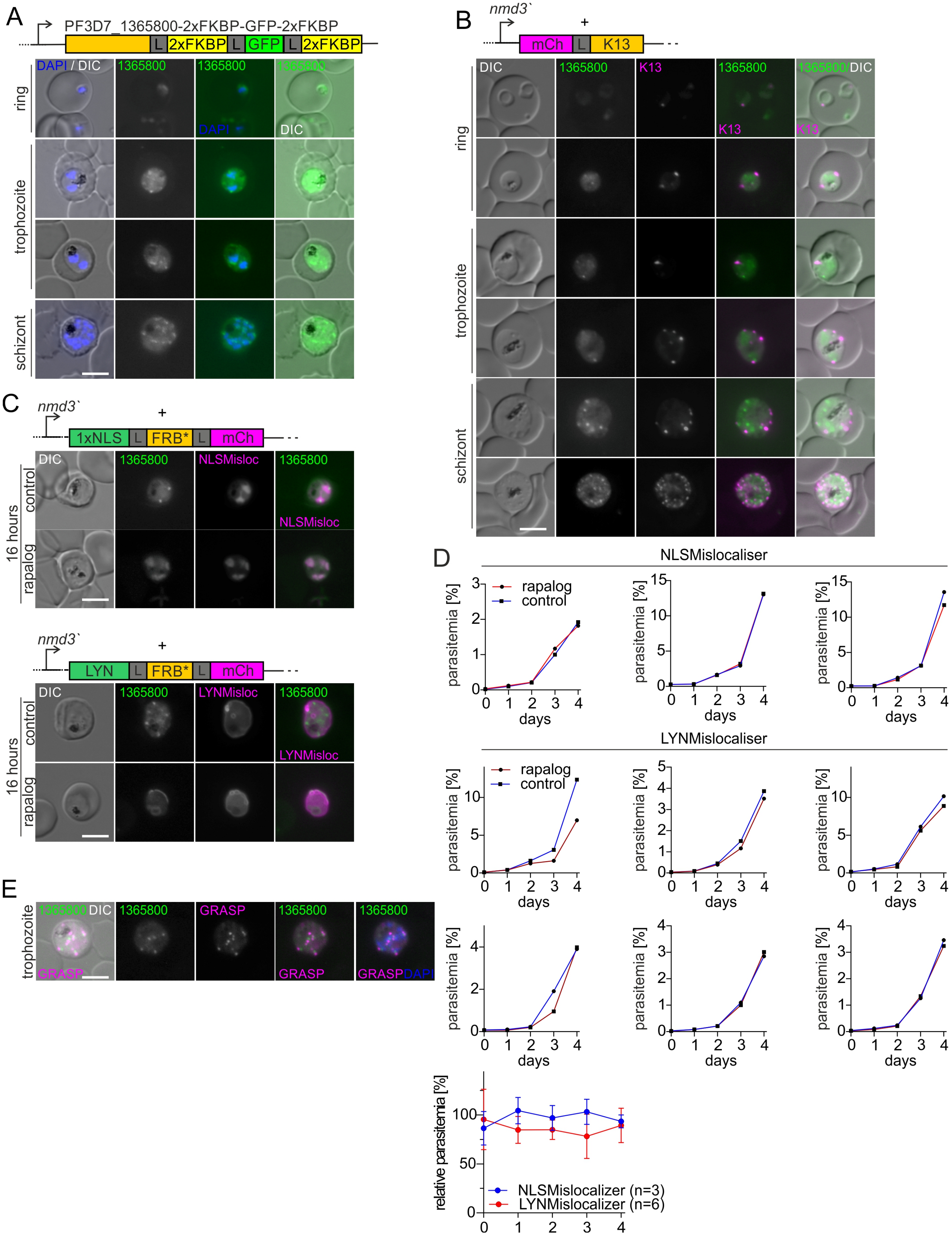

A
PF3D7_1365800-2xFKBP-GFP-2xFKBP
L 2xFKBP L GFP L 2xFKBP
DAPI / DIC
1365800
DAPI
DIC
ring
trophozoite
schizont
B
nmd3`
+
mCh L K13
DIC
1365800
K13
1365800/DIC
C
nmd3`
+
1xNLS L FRB* L mCh
NLSMisloc
16 hours
control
rapalog
LYN L FRB* L mCh
LYNMisloc
D
NLSMislocaliser
LYNMislocaliser
rapalog
control
parasitemia [%]
days
relative parasitemia [%]
NLSMislocalizer (n=3)
LYNMislocalizer (n=6)
E
1365800 DIC
GRASP
GRASP DAPI
trophozoite

### Figure S8

## A

| | Domain | Interpro | Protein |
|---|---|---|---|
| 1 | GAR<br>Binds to microtubules. | IPR003108 | KIC2 |
| 2 | PAH<br>Paired amphipathic helix, protein-protein interaction domain to act as a scaffold for transcription factors. | IPR036600 | KIC3 |
| 3 | VHS<br>Found at N-termini of proteins associated with endocytocis and vesicular trafficking. Might function as a multipurpose docking adapter that localizes proteins to the membrane through interactions with the membrane and/or the endocytic machinery | IPR002014 | KIC4<br>UBP1 |
| 4 | GAT<br>Found in eukaryotic GGAs (Golgi-localized gamma ear-containing Arf-binding proteins). Molecular anchor to trans-Golgi membrane via interaction with Arfs; interacts with rabaptin5 and ubiquitin. | IPR004152 | KIC4<br>KIC5 |
| 5 | Clathrin adaptor, α/β/γ-adaptin, appendage, Ig-like subdomain<br>Occurs in both major types of clathrin adaptors: AP complexes and GGAs. | IPR008152 | KIC4<br>KIC5 |
| 6 | β-adaptin appendage, C-terminal subdomain<br>Subdomain of the appendage (ear) domain of beta-adaptin. Required for binding to clathrin. A hydrophobic patch in the domain binds to a subset of D-phi-F/W motif-containing proteins (epsin, AP180, eps15). | IPR015151 | KIC4 |
| 7 | Clathrin adaptor, α-adaptin, appendage, C-terminal subdomain<br>Subdomain of the appendage (ear) domain of alpha-adaptin. The C-terminal appendage domain regulates translocation of endocytic accessory proteins to the bud site. | IPR003164 | KIC5 |

| | Domain | Interpro | Protein |
|---|---|---|---|
| 8 | ArfGAP<br>Occurs in GTPase activating proteins (GAPs) that activated the GTP hydrolysis activity of Arfs. Arfs participate in coat protein recruitment for membrane budding and fission. Before vesicles fuse with an acceptor compartment the membrane must be uncoated. This step requires the hydrolysis of Arf-associated GTP which is facilitated by an ArfGAP. The ArfGAP domain displays no obvious similarity to other GAP proteins. | IPR001164 | KIC7 |
| 9 | Metacaspase | IPR029030 | MCA2 |
| 10 | EH | IPR000261 | EPS15 |
| 11 | Ubiquitin specific protease domain (USP) | IPR028889 | UBP1 |
| 12 | Kelch-type beta propeller | IPR015915 | K13 |
| 13 | BTB/POZ domain | IPR000210 | K13 |
| 14 | Clathrin adaptor, µ subunit | IPR001392 | Ap-2µ |
| 15 | µ homology domain | IPR028565 | Ap-2µ |
| 16 | Myosin, N-terminal, SH3-like | IPR004009 | MyoF |
| 17 | Myosin head, motor domain | IPR001609 | MyoF |
| 18 | Ras-like-GTPase | cd00882 | MyoF |
| 19 | WD40 repeat | IPR001680 | MyoF |
| 20 | Tetratricopeptide-like helical domain superfamily (TPR) | IPR011990 | KIC11 |
| 21 | DnaJ | IPR001623 | KIC11 |

## B

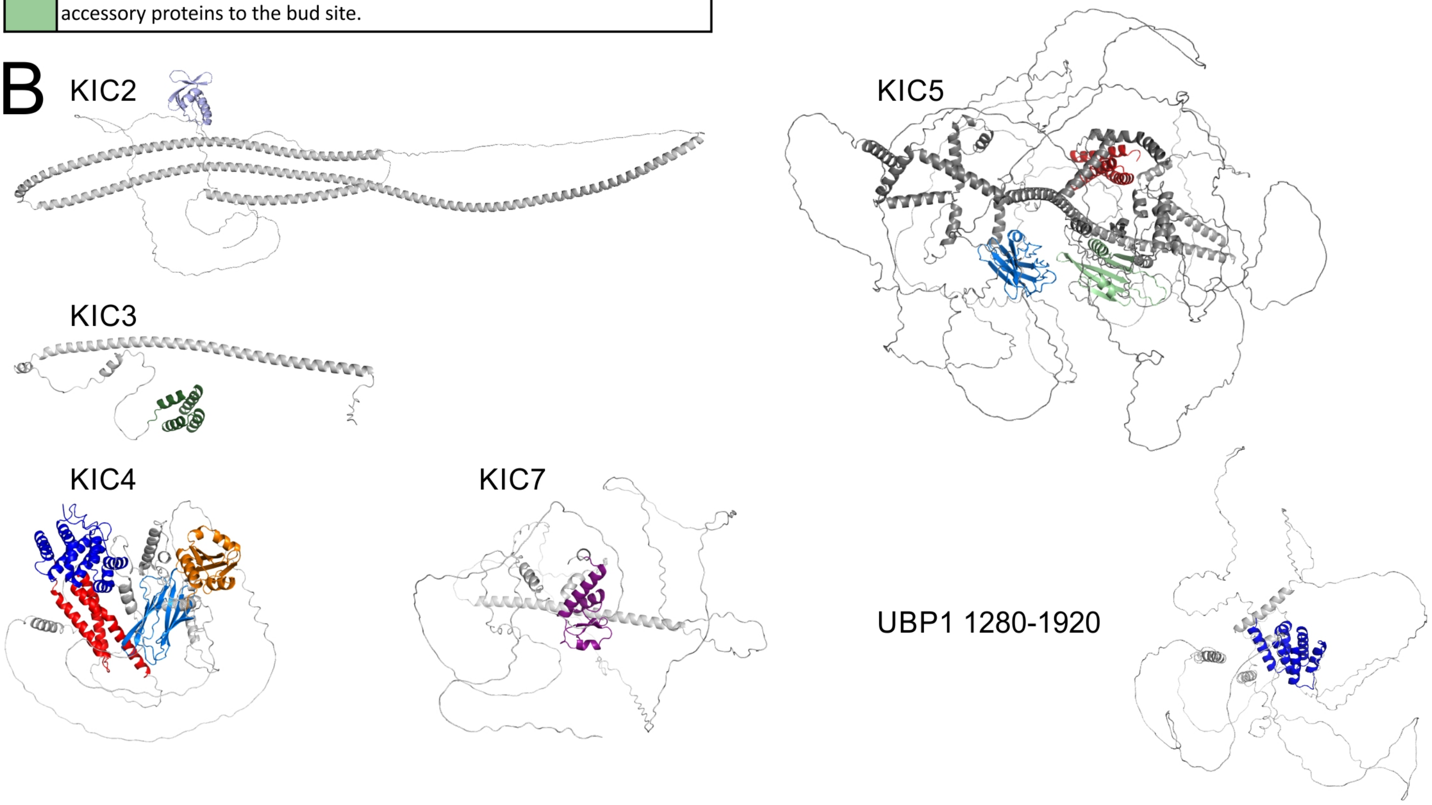


## C

PF3D7_1365800

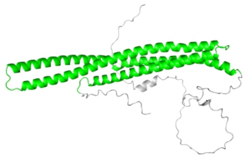

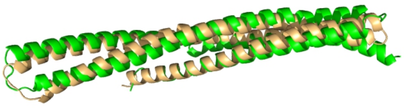

PF3D7_1365800 20-229 vs.
AH domain (4DCN)
RMSD 3.6 Å over 192 aa

AH domain IPR010504

Arfaptin homology (AH) domains dimerize and bind Arf and Rho family GTPases, including Arf1, which is invovled in vesicle budding and fusion. AH domains belong to the AH/BAR superfamily. BAR domains are required to tubulate membranes in early endocytosis.

### Figure S9

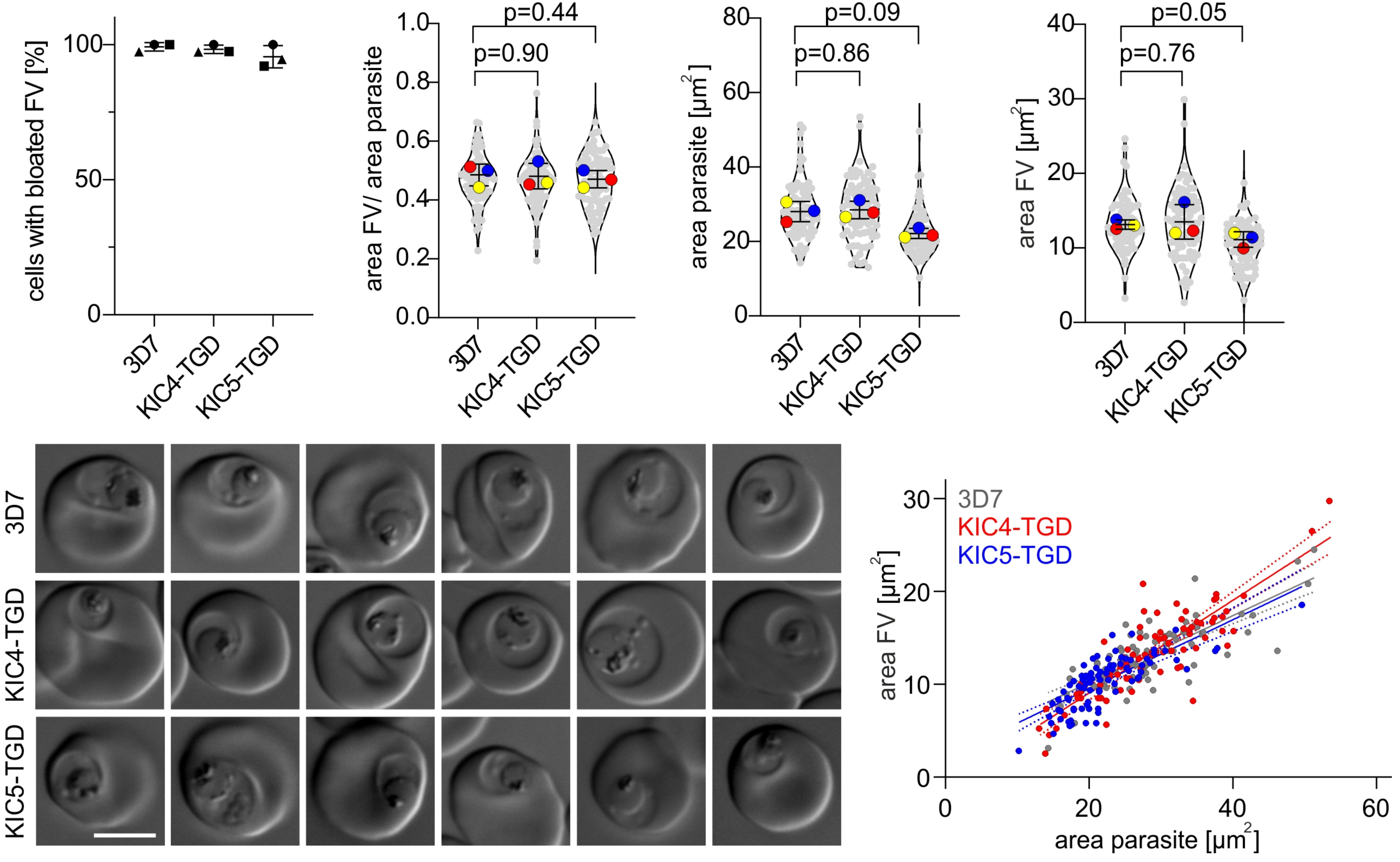

cells with bloated FV [%]
100
50
0
3D7
KIC4-TGD
KIC5-TGD
p=0.44
p=0.90
area FV/ area parasite
1.0
0.8
0.6
0.4
0.2
0.0
p=0.09
p=0.86
area parasite [µm2]
80
60
40
20
0
p=0.05
p=0.76
area FV [µm2]
40
30
20
10
0
3D7
KIC4-TGD
KIC5-TGD
area FV [µm2]
30
20
10
0
0
20
40
60
area parasite [µm2]

| Table S1 | | | | |
|---|---|---|---|---|
| **Gene ID** | **PlasmoDB annotation** | **Protein Name or Symbol** | **average log2 Ratio normalized Kelch13 from Birnbaum et al. 2020** | **proteins analyzed in this study** |
| PF3D7_1343700 | kelch protein K13 | K13 | 6.29 | |
| PF3D7_0606000 | protein KIC1 | KIC1 | 4.43 | |
| PF3D7_1227700 | protein KIC2 | KIC2 | 4.03 | |
| PF3D7_1246300 | protein KIC4 | KIC4 | 3.83 | |
| PF3D7_0914400 | protein KIC4 | KIC3 | 3.64 | |
| PF3D7_1438400 | metacaspase-like protein | MCA2 | 3.63 | X |
| PF3D7_1025000 | Eps15-like protein | Eps15 | 3.50 | |
| PF3D7_1138700 | protein KIC5 | KIC5 | 3.39 | |
| PF3D7_1243400 | conserved Plasmodium protein, unknown function | | 3.24 | X |
| PF3D7_0813000 | protein KIC7 | KIC7 | 3.13 | |
| PF3D7_1442400 | protein KIC9 | KIC9 | 3.03 | |
| PF3D7_0609700 | protein KIC6 | KIC6 | 2.98 | |
| PF3D7_0104300 | ubiquitin carboxyl-terminal hydrolase 1, putative | UBP1 | 2.90 | |
| *PF3D7_0204300* | *conserved Plasmodium protein, unknown function* | | 2.84 | |
| PF3D7_1014900 | protein KIC8 | KIC8 | 2.78 | |
| PF3D7_0915400 | ATP-dependent 6-phosphofructokinase | PFK9 | 2.58 | |
| *PF3D7_1365800* | *conserved Plasmodium protein, unknown function* | | 2.53 | X |
| PF3D7_1122500 | protein KIC10 | KIC10 | 2.44 | |
| PF3D7_1447800 | calponin homology domain-containing protein, putative | | 2.19 | X |
| PF3D7_1142100 | conserved Plasmodium protein, unknown function | KIC11* | 1.65 | X |
| PF3D7_1345600 | inner membrane complex protein | | 1.55 | |
| *PF3D7_0103100* | *vacuolar protein sorting-associated protein 51, putative* | *VPS51* | 1.55 | X |
| *PF3D7_0109000* | *photosensitized INA-labeled protein PHIL1, putative* | *PhIL1* | 1.49 | |
| *PF3D7_1117900* | *conserved Plasmodium protein, unknown function* | | 1.46 | |
| PF3D7_0408100 | conserved Plasmodium protein, unknown function | | 1.45 | |
| PF3D7_1247400 | peptidyl-prolyl cis-trans isomerase FKBP35 | FKBP35 | 1.45 | |
| *PF3D7_0721100* | *conserved Plasmodium protein, unknown function* | | 1.42 | |
| *PF3D7_1016200* | *Rab3 GTPase-activating protein non-catalytic subunit, putative* | | 1.41 | |
| PF3D7_0708400 | heat shock protein 90 | HSP90 | 1.41 | |
| PF3D7_1329100 | myosin F | MyoF | 1.32 | X |
| PF3D7_1329500 | conserved Plasmodium protein, unknown function | KIC12* | 1.32 | X |
| PF3D7_0717600 | conserved Plasmodium protein, unknown function | IMC32 | 1.30 | |
| PF3D7_0405700 | lysine decarboxylase, putative | UIS14 | 1.23 | X |
| *PF3D7_0907200* | *GTPase-activating protein, putative* | | 1.15 | X |
| *PF3D7_0822900* | *conserved Plasmodium protein, unknown function* | *PIC2* | 1.15 | |
| *PF3D7_1018200* | *serine/threonine protein phosphatase 8, putative* | *PPP8* | 1.13 | |
| PF3D7_1141300 | conserved Plasmodium protein, unknown function | APR1 | 1.11 | |

| | |
|---|---|
| | KICs (Birnbaum et al. 2020) |
| | new candidates |
| | new candidates not analyzed |
| | BioID baits (Birnbaum et al. 2020) |
| | considered false positives / IMC proteins |
| * | renamed in current manuscript |
| *italic* | less stringent filtering group (significant with FDR<1% in 2 out of 4 reactions of any bait (Birnbaum et al. 2020)) |

| Table S2: Oligonucleotides | |
|---|---|
| **name** | **sequence** |
| MCA2(wt)-3xHA-T2A-Neo fw | ggtgacactatagaatactcaagctgcggccgcTAATGAGTACTCACCAGATGACATCAAATTTTTATCAC |
| MCA2(wt)-3xHA-T2A-Neo rv | CTGGAACATCGTATGGGTACATGGTGGTACCGGAAACACATTTAATATTCAAATCGATAATACC |
| MCA2(wt)-3xHA int check fw | GGAATATAATTTGAAATATGTAAAATTTAAACTGTCC |
| MCA2(wt)-3xHA int check rv | CATAAACACAAAAATTTAAAGATGGAGTGG |
| MCA2 (Y1334.)-GFP fwd | ggtgacactatagaatactcgcggccgctaaAATAATTTTAGCAAACCAAATTTTTTAGATAAATTTTTTTATG |
| MCA2 (Y1334.)-GFP rev | CAGCACCAGCAGCAGCACCTCTAGCacgcgtTTTTTTTAATTGTTCATATAACTTTTTATTTTTGGTC |
| Intcheck 5UTR MCA2 (Y1344.) fw | ATCCATAACAATAATAATAATATTGAGTGG |
| Intcheck 3UTR MCA2 (Y1344.) rv | AACTTTTTTGTTTTGGTATCTTTGTCAAGC |
| PfMyosinC_NotI_fwd | cagtgcggccgctaaatagtaataaaacatatgatgaaag |
| PfMyosinC_AvrII_rev | cagtcctaggtacaaaagacctgcgccagaaaac |
| Intcheck 5UTR MyoF wt fw | CAAATTATCAGGTCATAAAAAGGCAATAACATG |
| Intcheck 3UTR MyoF wt rv | ATATATATATATATGAACAAAATTTACAATATC |
| MyoF intcheck fw JSW175 | GGCACACTTAAATCATATGAAC |
| MyoF intcheck rv JSW176 | catgtgaatagaaaagtaaac |
| MyoF(wt)-3xHA-T2A-Neo fw | gctatttaggtgacactatagaatactcaagctgcggccgcTAATCTCAGAAGATAAAATTATTTCATCATC |
| MyoF(wt)-3xHA-T2A-Neo rv | CGTAATCTGGAACATCGTATGGGTACATGGTGGTACCTACAAAAGACCTGCGCCAGAAAACTATG |
| KIC11 TGD fw | gctatttaggtgacactatagaatactcgcggccgctaaAACGATAAAAAGAATAGCATTAATAAAAG |
| KIC11 TGD rv | cagcaccagcagcagcacctctagcacgcgtTATATGATTATTCTTTTTATTTATAG |
| KIC11 fw | gctatttaggtgacactatagaatactcgcggccgctaaGATGATGCTGACGAGGAGGAAGAAG |
| KIC11 rv | CAGCAGATCTTGATCTCAATCCTGAcctaggTTTTTTATCCTTTGTTTTAAGTTTAT |
| KIC11 int check 5 fw | ATGTAACATATGATAATAATAATG |
| KIC11 int check 3 rv | TATTAAAAAGAATTTCATTTGATG |
| KIC12fw | gctatttaggtgacactatagaatactcgcggccgctaaAGAGTGGTGGTAGTAACAACAATAG |
| KIC12 rv | CAGCAGATCTTGATCTCAATCCTGAcctaggATAATTTAACTCTGGGTGACCACTAAAC |
| KIC12 TGD fw | gctatttaggtgacactatagaatactcgcggccgctaaGGACAATTTATATTATAATAAATATAC |
| KIC12 TGD rv | cagcaccagcagcagcacctctagcacgcgtCATGTTTTTCTTGGCATTATAAAATG |
| KIC12 int check 5 fw | TTACAGCATATGTAAAGAATTCATC |
| KIC12 int check 3 rv | TTTCCAATCAACCTATATATGTGTG |
| KIC12 TGD int check 5 fw | AATGGGCATATTCTATTTTCGTGTG |
| KIC12 TGD int check 3 rv | AACATCAAGCAAAAAATAACATCC |
| VPS51 fw | tatttaggtgacactatagaatactcgcggccgctaaAGTGATATGTATGAAAAAATTGAAG |
| VPS51 rv | AGCAGCAGATCTTGATCTCAATCCTGAcctaggCTCTTTAAACATTTTCAATATAAATAATTTATTTTC |
| VPS51 int check 5 fw | TGAAGGACAGGAAAGAGAATATG |
| VPS 51 int check 3 rv | ATGAATCTATCCTTTCTTTACTC |
| VPS51 TGD fw | gctatttaggtgacactatagaatactcgcggccgctaaAATAAAAATAATAGAAGAAAAAATG |
| VPS51 TGD rv | ccagcaccagcagcagcacctctagcacgcgtCTTTTTCGATTTATTTATTAAATC |
| VPS51 TGD int check 5 fw NEW | cgcctgggagtcattcctaaatttg |
| VPS 51 TGD int check 3 rv NEW | GATGTACACATTTGTTATTATCCC |
| PF3D7_1243400 rv | AGCAGCAGCAGATCTTGATCTCAATCCTGAcctaggAGAAGCTACCTTTTGTATTTCC |
| PF3D7_1243400fw | aagctatttaggtgacactatagaatactcgcggccgctaaGAGAACGAACAAAAGATATCCTTC |
| PF3D7_1447800 fw | gctatttaggtgacactatagaatactcgcggccgctaaAATAATATACATAATAATAATAAAC |
| PF3D7_1447800 rv | CAGCAGATCTTGATCTCAATCCTGAcctaggATCTTGACCTGTGATCATATTTTC |
| PF3D7_1447800 int check 5 fw | ATGATAAAAATAATAATGCACATAG |
| PF3D7_1447800check 3 rv | AGCAATATATCTAACCAATGGACAC |
| PF3D7_1365800 fw | gctatttaggtgacactatagaatactcgcggccgctaatattttagAAATTCAAGTATCAGG |
| PF3D7_1365800 rv | CAGCAGATCTTGATCTCAATCCTGAcctaggTTGGGGGAATATAGGTGTAATATTAG |
| PF3D7_1365800 int check 5 fw | tatatgtgcacatatttatatttac |
| PF3D7_1365800 int check rv NEW JSW | gtgatgtccatataaataatgttgac |
| PfUIS14fw | gctatttaggtgacactatagaatactcgcggccgctaaATTGTAATTATGTTAAAAAATGTG |
| PfUIS14 rv | CAGCAGATCTTGATCTCAATCCTGAcctaggCTTGTGTTTGGTTTCACAGTCATC |
| PfUIS14 TGD fw | gctatttaggtgacactatagaatactcgcggccgctaaGAACAATATAGATCAGAATAAAATC |
| PfUIS14 TGD rv | cagcaccagcagcagcacctctagcacgcgtATTAATTATCTCATGTAAATTTTCTG |
| PfUIS14 int check 5 fw | CTAGCTAGTAATAACAATTATTGTG |
| PfUIS14 int check 3 rv | ATATAATATATATAATTCCATGCTG |
| PfUIS14 TGD int check 5 fw | TTTCAAATCGATGAGGATTCTTTAC |
| PfUIS14 TGD int check 3 rv | TAGAAACTTCTTTATATTTTTTATC |
| PF3D7_0907200 fw | gctatttaggtgacactatagaatactcgcggccgctaaTGCATGAACTTAATATGTGTGATAATG |
| PF3D7_0907200 rv | CAGCAGATCTTGATCTCAATCCTGAcctaggAAATGTGTGAGCATCGTCTACCAG |
| PF3D7_0907200 int check 5 fw | TAATCAAAATATAAAAAATCATCAC |
| PF3D7_0907200 int check 3 rv | TCAAGAAAAAAAATTTAATTATACG |
| PF3D7_1243400 TGD fw | cactatagaatactcgcggccgctaaTTTCCCAGTTTTAATAA |
| PF3D7_1243400 TGD rv | ctgatattaacttctgctcgtttaaacCTTCTCTTGGTTATCTTGTA |
| ama1 fw NotI | GGTGACACTATAGAATACTCgcggccgcGAGGTGTGTTGGGAAACAGAAG |
| ama1 rv KpnI | TCCTggtaccTTTGTACAATTTATAACAAGTAC |
| IMC1c fw KpnI | CTCGggtaccATGGCAGATTCAATCAAAAGTTCAAACAG |
| ARO fw KpnI | CTCGggtaccATGGGAAATAATTGCTGTGCAGGAAG |
| AMA1 fw KpnI | CTCGggtaccATGAGAAAATTATACTGCGTATTATTATTG |
| mCherry rv XmaI | GAACATTAAGCTGCCATATCCCTCGACCCGGGTTACTTGTACAGCTCGTCCATGCCGCC |
